# Myricetin allosterically inhibits Dengue NS2B-NS3 protease as studied by NMR and MD simulations

**DOI:** 10.1101/2021.12.13.472523

**Authors:** Mei Dang, Liangzhong Lim, Amrita Roy, Jianxing Song

## Abstract

Dengue NS2B-NS3 protease existing in equilibrium between the active and inactive forms is essential for virus replication, thus representing a key drug target. Here Myricetin, a plant flavonoid, was characterized to non-competitively inhibit Dengue protease. Further NMR study identified the protease residues perturbed by binding to Myricetin, which were utilized to construct the Myricetin-protease complexes. Strikingly, in the active form Myricetin binds a new allosteric site (AS2) far away from the active site pocket and allosteric site (AS1) for binding Curcumin, while in the inactive form it binds both AS1 and AS2. To decipher the mechanism for the allosteric inhibition by Myricetin, we conducted molecular dynamics (MD) simulations on different forms of Dengue NS2B-NS3 protease. Unexpectedly, the binding of Myricetin to AS2 is sufficient to disrupt the active conformation by displacing the characteristic NS2B C-terminal β- hairpin from the active site pocket. By contrast, the binding of Myricetin to AS1 and AS2 results in locking the inactive conformation. Therefore Myricetin represents the first small molecule which allosterically inhibits Dengue protease by both disrupting the active conformation and locking the inactive conformation. The results enforce the notion that a global allosteric network exists in Dengue NS2B-NS3 protease, which is susceptible to allosteric inhibition by small molecules such as Myricetin and Curcumin. As Myricetin has been extensively used as a food additive, it might be directly utilized to fight the Dengue infections and as a promising starting for further design of potent allosteric inhibitors.

**Graphic Abstract:** 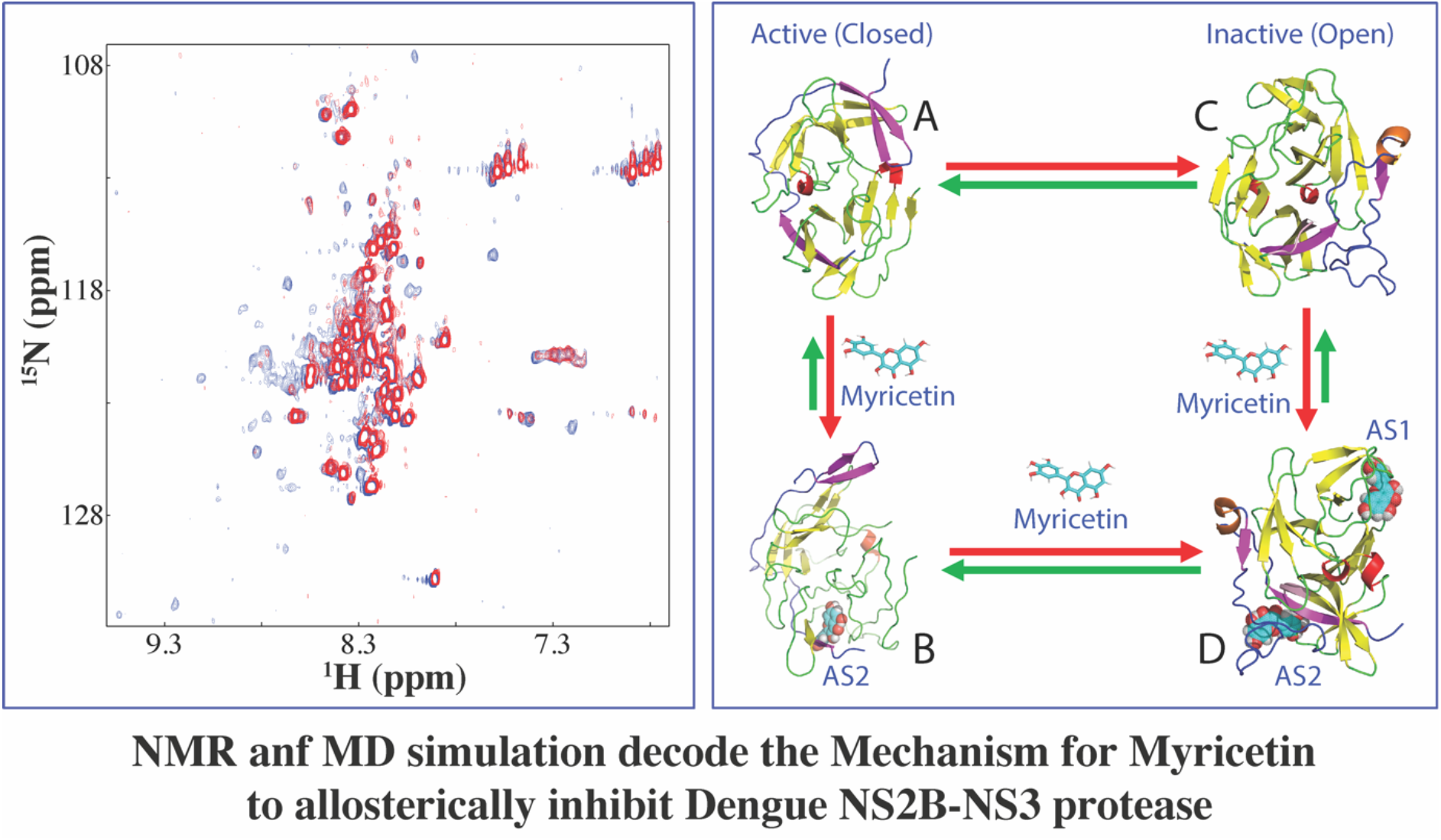

## 1. INTRODUCTION

Dengue virus (DENV) belongs to the *Flaviviridae* family, which also includes Zika (ZIKV), West Nile (WNV), Japanese encephalitis and Yellow fever viruses. DENV is the most prevalent human pathogens transmitted by *Aedes* mosquitoes with 3.6 billion people at risk in >120 countries, particularly in tropical and subtropical regions such as Singapore (1–3). It still remains a large challenge to develop effective vaccine and so far no marketed antiviral drug exists to treat dengue associated diseases despite exhaustive studies (4, 5).

Common to all *Flaviviridae* members, DENV has an ∼11-kb single-stranded positive sense RNA genome, which is translated into a large polyprotein by the host-cell machinery. Intriguingly, the large polyprotein needs to be subsequently processed into 10 proteins, which include three structural (capsid, membrane, and envelope) and seven nonstructural (NS1, NS2A/B, NS3, NS4A/B, and NS5) proteins. The processing of the polyprotein is conducted by the host cell proteases such as furin and signalaseas, as well as a virus-encoded protease, which thus represents an established target for drug design to treat DENV and other flavivirus infections (4–26).

The Dengue protease domain is over the N-terminal part of NS3 protein, which folds into a chymotrypsin-like fold consisting of two β-barrels with each composed of six β-strands, and the catalytic triad (His51-Asp75-Ser135) located at the cleft between the two β-barrels. Intriguingly, unlike other chymotrypsin-like proteases such as 3C-like proteases of coronaviruses (23), the flavivirus proteases including Dengue one require an additional stretch of ∼40 amino acids from the cytosolic domain of NS2B protein for its correct folding and catalytic function, thus called a two-component NS2B-NS3 protease (7–24). In crystal structures of flaviviral NS2B-NS3 proteases determined so far, the NS3 protease domains adopt highly similar structures, while the NS2B cofactor was found to assume two distinctive conformations: namely the active or closed form upon being complexed with the substrate peptide as well as the inactive or open form in the free state (Fig. S1). Noticeably, in the active form, the NS2B cofactor becomes wrapped around the NS3 protease domain with its C-terminal residues Ser75-Ser79 and Gly82-Ile86 forming a short β-hairpin to serve as part of the active site pocket (7–10). Furthermore, NMR studies revealed that in solution the two forms undergo significant exchanges, thus leading to a high conformational dynamics on μs-ms time scale (12–15). Strikingly, it appears that a global allosteric network exists in Dengue NS2B- NS3 protease and the perturbation of key residues/sites of the network is sufficient to modulate the conformational equilibrium, thus manifesting as the allosteric effect. For example, the chemical modification of residues around Ala125 led to the conformational equilibrium being shifted to and then locked in the inactive form (Fig. S1), which has both NS2B cofactor and NS3 protease domain well-defined in the crystal structure (17). Furthermore, as we recently showed by NMR and MD simulations, the binding of a natural product Curcumin to a pocket close to Ala125 without any overlap with the active site resulted in the allosteric inhibition of Dengue protease by destabilizing the active conformation (26).

To date, tremendous efforts have been dedicated to drug development by targeting flaviviral NS2B-NS3 proteases but the results revealed the major challenge in rational design of their active site inhibitors: their active sites are relatively flat (8–26). So one promising strategy to overcome this challenge is to discover/design their allosteric inhibitors. Previously, to respond to the urgency to fight ZIKV and DENV infection in Singapore, we have conducted intense efforts to screen inhibitors for Zika and Dengue NS2B-NS3 proteases from natural products isolated from edible plants. As a result, we have successfully identified Curcumin and a group of flavonoids with significant inhibitory effects on both proteases, which include Myricetin, Quercetin, Luteolin, Isorhamnetin and Apigenin, out of which Myricetin inhibits Zika NS2B-NS3 protease with the highest activity (IC50 of 1.3 uM and Ki of 0.8 uM) in a non-competitive mode. In fact, plant flavonoids have been previously shown to inhibit Dengue protease (24, 25), but so far their binding sites and inhibitory mechanisms have not been experimentally characterized such as by X-ray crystallography or NMR spectroscopy, most likely due to the challenge in studying Dengue NS2B-NS3 protease with significantly- provoked dynamics upon binding to these flavonoids.

Recently, with a protocol we previously developed for selectively isotope-labelling the NS2B cofactor or NS3 protease domain (15), we have successfully utilized NMR spectroscopy to identify the binding-involved residues by Curcumin, although the binding by Curcumin did dramatically enhance the conformational changes on μs-ms time scale (26). Consequently, we were able to construct the structures of the Curcumin-protease complexes with NMR-derived constraints and conducted further molecular dynamics (MD) simulations. The results revealed that despite binding to a pocket without any overlap with the active site, Curcumin achieves the allosteric inhibition by disrupting the active conformation of Dengue protease (26).

Here we aimed to decode the mechanism by which Myricetin inhibits Dengue protease by enzymatic assay, NMR characterization, molecular docking and MD simulations. Briefly, the enzymatic assay showed that Myricetin inhibits Dengue protease also in a non-competitive manner. Further NMR studies identified the binding-perturbed residues by Myricetin, which are dramatically different from those by binding to Curcumin (26). With NMR-derived constraints, the structures of the Myricetin-protease complexes have been successfully constructed for both active and inactive forms. Very unexpectedly, in the active form, Myricetin binds to a new allosteric site (AS2) far away from the allosteric site (AS1), which was previously identified for binding Curcumin (Fig. S1B) (26), By contrast, in the inactive form, Myricetin binds to both AS1 and AS2. Subsequent MD simulations decode that the Myricetin-binding to the active form at AS2 far away from the active site pocket is sufficient to allosterically destabilize the active conformation. By contrast, the Myricetin-binding to the inactive form at both AS1 and AS2 locks the inactive conformation. The results together indicate that Dengue protease has more than one allosteric sites for natural products and is susceptible to the allosteric inhibition. Therefore allosteric inhibitors could be designed to target Dengue NS2B-NS3 protease not only by disrupting the active conformation but also by locking the inactive conformation.

## 2. RESULTS

### 2.1. Myricetin inhibits Dengue NS2B-NS3 protease in a non-competitive manner

Previously to obtain the active recombinant Dengue NS2B-NS3 protease in *E. coli* cells, the enzyme was extensively constructed by joining the NS2B cofactor and NS3 protease domain covalently with an engineered linker. However, this artificial form does not exist *in vivo* and in particular has NMR spectra of poor quality (12–15). Therefore to solve this problem, we have previously developed a protocol to recombinantly generate Dengue protease with the NS2B cofactor and NS3 protease domain unlinked (15), which also allowed to selectively isotope-label either NS2B or NS3 in the Dengue NS2B-NS3 protease complex for detailed NMR studies as demonstrated in our recent delineation of the mechanism for the allosteric inhibition by Curcumin (26).

In the present study, we used the same unlinked Dengue protease for all enzymatic and NMR experiments. We have determined its Km to be 89.39 ± 6.62 μM; and Kcat to be 0.12 ± 0.01 s^-1^, which are almost identical to our previous results with Km of 92.39 ± 9.94 μM and Kcat of 0.15 ± 0.01 s^-1^ (15, 26). In our preliminary screening (20), we found that Myricetin showed significant inhibition on both Zika and Dengue NS2B-NS3 proteases. Here, we further determined its IC50 to be 8.46 ± 0.48 μM and inhibitory constant Ki to be 4.92 ± 0.21 μM on Dengue protease (Fig. 1A), which are very similar to those of Curcumin on Dengue protease (IC50 of 7.18 and Ki of 4.35 μM) (26). Intriguingly, the inhibitory activity of Myricetin on Dengue protease appears to be lower than those on Zika protease (IC50 of 1.30 μM and Ki of 0.80 μM) (20), which might be due to distinctive physicochemical properties, conformations or/and dynamics of two proteases. However, similar to what we previously observed on Zika protease (20), Myricetin also inhibited Dengue protease by changing Vmax but not Km (Fig. 1A), thus suggesting that Myricetin acts as a non-competitive inhibitor for Dengue protease.

**Figure 1.**
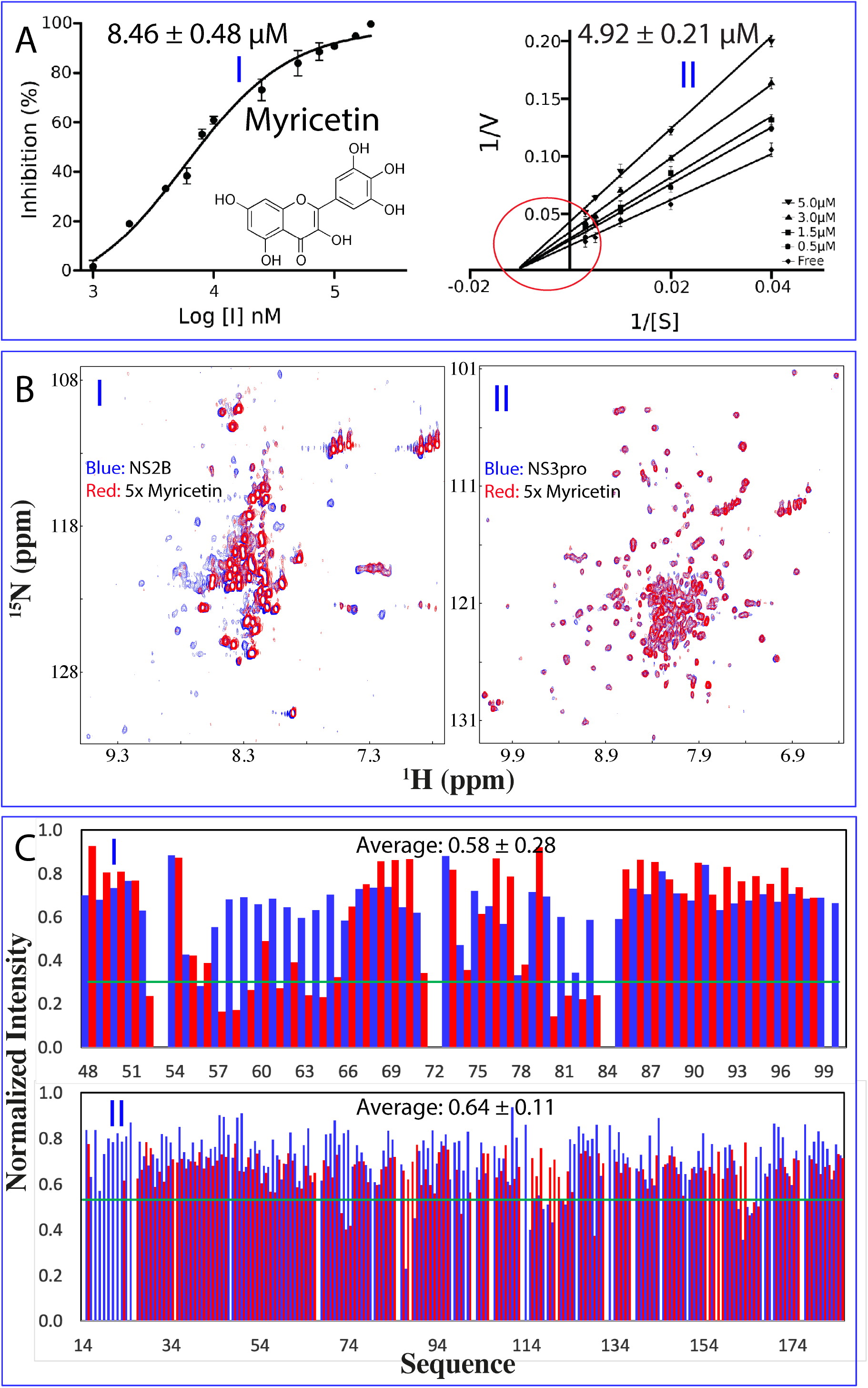
Enzymatic and NMR characterization of the inhibition of Dengue NS2B-NS3 protease by Myricetin. (A) Chemical structure of Myricetin and the inhibitory data used for fitting IC50 for Myricetin. Lineweaver-Burk plot for determining inhibitory constant (Ki) of Myricetin. [S] is the substrate concentration; v is the initial reaction rate. The red circle is used to indicate that the inhibition is non-competitive characteristic of the same Km but varying Vmax values. (B) (I) NMR ^1^H- ^15^N HSQC spectra of the Dengue NS2B-NS3 complex with NS2B selectively ^15^N-labeled at a protein concentration of 100 µM in the absence (blue) and in the presence of Myricetin (red) at 1:5 (Proteae:Myricetin). (II) HSQC spectra of the protease complex with NS3 selectively ^15^N-labeled at a protein concentration of 100 µM in the absence (blue) and in the presence of Myricetin (red) at 1:5. (C) (I) Normalized HSQC peak intensity of the ^15^N-labeled NS2B complexed with the unlabeled NS3 in the presence of Myricetin at 1:5. (II) Normalized HSQC peak intensity of the ^15^N-labeled NS3 complexed with the unlabeled NS2B in the presence of Myricetin at 1:5. Significantly perturbed residues are defined as those with the normalized intensity < 0.53 for NS2B and < 0.59 for NS3 (average value - one standard deviation).

### 2.2. NMR characterization of the binding of Myricetin to Dengue NS2B-NS3 protease

To determine the binding modes of Myricetin, here we prepared the Dengue protease samples with either NS2B cofactor or NS3 protease domain selectively ^15^N-labeled. As shown in Fig. 1B, in the unlinked Dengue protease complex, both ^15^N-labeled NS2B (I of Fig. 1B) and NS3 (II of Fig. 1B) have well-dispersed HSQC spectra, indicating that both of them are well-folded in the protease complex. In particular, the chemical shifts of their HSQC peaks are very similar to what were previously reported (13,15,26).

We then titrated the Dengue protease samples with Myricetin at molar ratios of 1:0.5, 1:1; 1:2.5, 1:5 and 1:10 (Protease:Myricetin). Intriguingly, no significant shifts were observed for most HSQC peaks of NS2B (I of Fig. 1C) and NS3 (II of Fig. 1C), indicating no major structural change upon binding, which is very similar to what was observed on the binding of Curcumin to Dengue protease (26). On the other hand, their HSQC peaks became stepwise broadened and consequently the intensity of the peaks gradually reduced. At 1:10, many well-dispersed HSQC peaks became too weak to be detectable. Usually, the line-broadening of HSQC peaks upon binding results from the micro-molar dissociation constants, and/or binding-induced increase of conformational exchanges particularly on µs-ms time scale (11-15,26-28). Indeed, for a folded but dynamic protein, a slight disruption/destabilization of the native structure is sufficient to trigger the significant increase of conformational exchange on µs-ms time scale, and consequently resulting in the broadening/disappearance of many well-dispersed HSQC peaks (28–30). Here, the Myricetin binding-provoked increase of conformational exchanges as detected by NMR might explain the results that our ITC measurements of the binding of both Myricetin and Curcumin (26) to Dengue protease all gave rise to the data with a very high level of noises.

The current results indicate that as we recently observed on Curcumin (26), the binding of Myricetin also led to significant increase of conformational dynamics particularly on µs-ms time scale, in a sharp contrast to a recent NMR study with an active-site inhibitor of Dengue protease, in which the inhibitor binding led to a significantly reduced dynamics, thus showing much better quality of NMR spectra (14). As such, despite intense attempts, the significant weakening of the intensity of HSQC peaks prevented us from acquiring high-quality NMR relaxation data to derive their backbone dynamics of the protease bound to Curcumin or Myricetin as we previously performed on other proteins to gain insights into the ps-ns and μs- ms dynamics (27, 28)

### 2.3. Distinctive binding modes of Myricetin for the active and inactive forms

Due to the significant increase of the dynamics upon binding to Myricetin or Curcumin (26), which led to severe NMR line broadening, we were also unable to further determine the structures of Dengue protease in complex with Myricetin or Curcumin by NMR spectroscopy. Furthermore, we also intensely attempted to crystalize their complex samples by screening a large array of buffer conditions but all failed. Nevertheless, the ability to selectively label the NS2B cofactor and NS3 protease domain offered us to follow the intensity changes of HSQC peaks upon adding Myricetin at different molar ratios.

Fig. 1C presents the normalized peak intensities of NS2B (I of Fig. 1C) and NS3 (II of Fig. 1C) in the presence of Myricetin at 1:5 (red bars) as well as Curcumin at 1:5 (blue bars). The NS2B residues have the average intensity of 0.58 and 0.63 for their HSQC peaks, while the NS3 residues have the average intensity of 0.64 and 0.76 in the presence of Myricetin and Curcumin respectively, implying that both Myricetin and Curcumin similarly triggered slightly higher dynamics for the NS2B cofactor than for the NS3 protease domain. Very unexpectedly, however, at residue-specific resolution, Myricetin and Curcumin induced the distinctive patterns of the intensity changes for both NS2B and NS3 residues. For example, for the NS2B cofactor, Myricetin additionally triggered the significant reduction (< average – one STD) of the HSQC peak intensity of the residues over 56-64 and 80-83 (I of Fig. 1C), while for the NS3 protease domain, Myricetin induced the complete disappearance of HSQC peaks of the residue over 18-23 (II of Fig. 1C).

We thus mapped these significantly-perturbed residues back to the Dengue protease structures in either active (Fig. 2A) or inactive (Fig. 3A) forms. With these NMR-derived constraints, we were able to construct the structures of the Myricetin-protease complexes for the active (Fig. 2B) or inactive (Fig. 3B) form by the well-established molecular docking program HADDOCK (31), which we extensively utilized to build up the protease-Curcumin complex (26) and complexes of other proteins with small molecules including ATP (32, 33).

**Figure 2.**
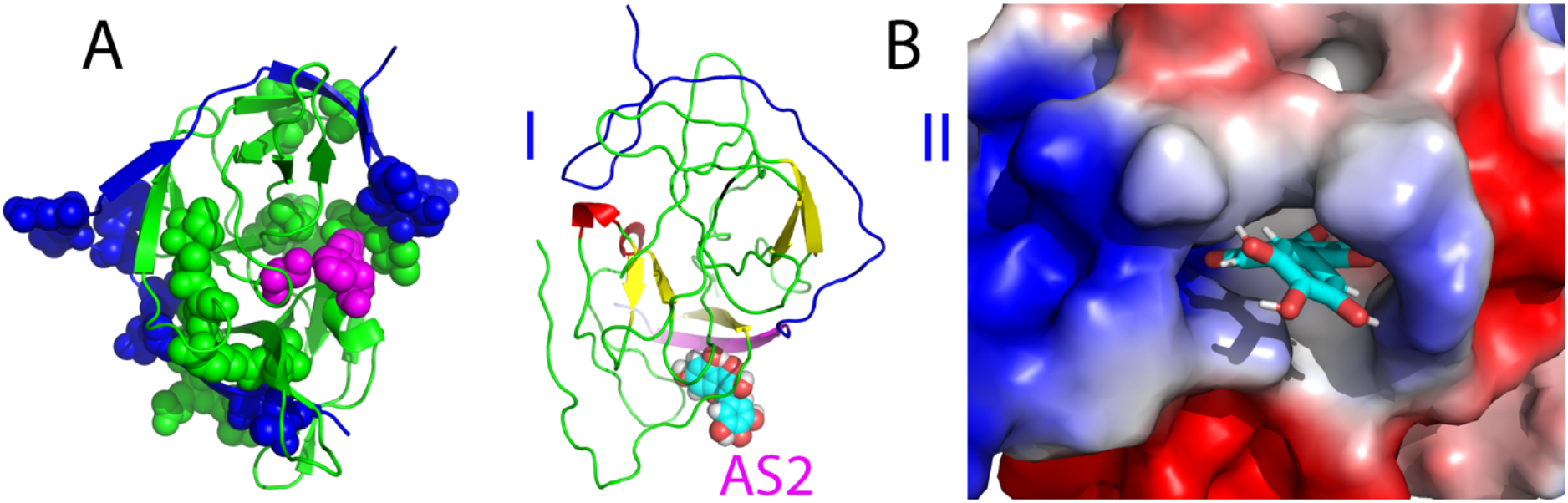
The active form of Dengue NS2B-NS3 protease complexed with Myricetin. (A) Structure of the active form of Dengue protease with the significantly perturbed residues displayed in spheres. NS2B is colored in blue while NS3 in green. The catalytic triad residues His51-Asp75-Ser135 are displayed in purple spheres. (B) The lowest-energy docking structures in ribbon (I) and electrostatic potential surface (II) of the active form of the protease complexed with Myricetin at a new allosteric site (AS1). The β-strand is colored in purple and loop in blue for NS2B, while the β-strand is in yellow, helix in red and loop in green for NS3. Myricetin is displayed in spheres (I) and sticks (II) respectively.

**Figure 3.**
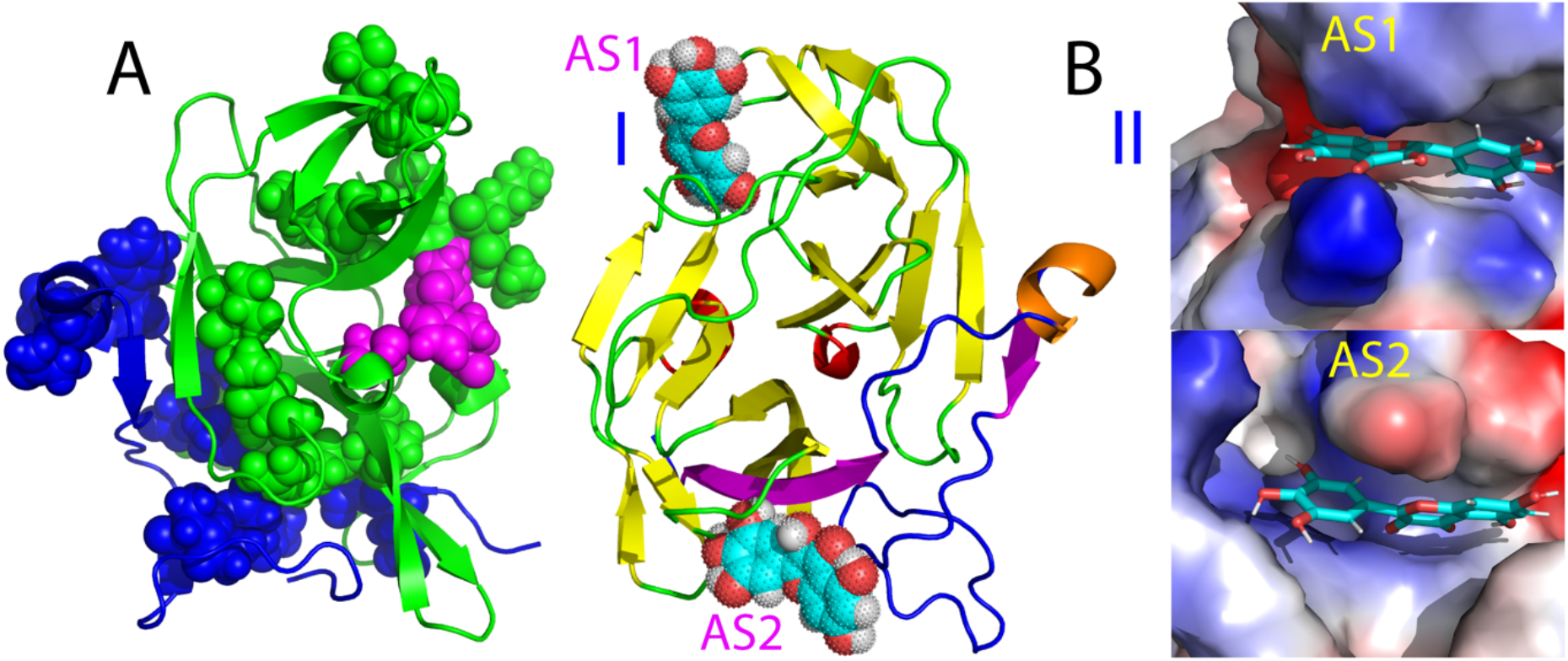
The inactive form of Dengue NS2B-NS3 protease complexed with Myricetin. (A) Structure of the inactive form of Dengue NS2B-NS3 protease with the significantly perturbed residues displayed in spheres. NS2B is colored in blue while NS3 in green. The catalytic triad residues His51-Asp75-Ser135 are displayed in purple spheres. (B) The lowest-energy docking structures in ribbon (I) and electrostatic potential surface (II) of the inactive form of the protease complexed with Myricetin at two allosteric sites (AS1 and AS2). The β- strand is colored in purple, helix in brown and loop in blue for NS2B, while the β-strand is in yellow, helix in red and loop in green for NS3. Myricetin is displayed respectively in spheres (I) and sticks (II).

Fig. 2B presents the lowest-energy structure of the Myricetin-protease complex for the active form. Very unexpectedly, completely different from the Curcumin-protease complex for the active form we previously obtained (Fig. S1B), in which Curcumin binds Dengue protease at an allosteric site (designated as AS1) close to Ala125 (17, 26), here Myricetin binds Dengue protease at a new allosteric site (designated as AS2) which is far away from both the active site pocket and AS1. It is very interesting to note that even in the complex, the short β-hairpin characteristic of the active form over the NS2B residues Ser75-Ser79 and Gly82-Ile86 is distorted to some degree although it is still located at the original position, implying that the binding of Myricetin to AS2 could perturb the conformation of this β-hairpin despite being located far away. A close examination revealed that AS2 is constituted mostly by the NS3 residues Lys15-Asp20 in loop and Gly21-Arg24 in β-stand, as well as the NS2B residues Leu54-Val59 (I of Fig. 2B). Consistent with the complex structure, in the presence of Myricetin at 1:5, HSQC peaks of many of these involved NS3 residues became too broad to be detectable while those of the NS2B residues also have significantly reduced intensities (Fig. 1C), both of which are unique for the Myricetin binding and were not observed on the Curcumin binding (Fig. 1C). Moreover, the AS2 is relatively positively charged as shown in II of Fig. 2B.

On the other hand, for the inactive form of Dengue protease (17), Myricetin was identified to bind not only AS2, but also AS1 we previously found for binding Curcumin. For AS2 of the inactive form, in addition to the NS3 and NS2B residues observed in the active form (Fig. 2B), the NS2B residues Thr77-Gly82 further fold back to have the direct contacts with Myricetin (Fig. 3B). For AS1 of the inactive form, only NS3 residues are involved in contacting Myricetin, including Val72-Asp75 and T118-T122 as well as G151-V154 which form a β-sheet with A160-A166. Again consistent with the complex structure, many of these involved NS3 residues have significantly reduced intensities of their HSQC peaks upon binding to Myricetin but not to Curcumin (II of Fig. 1C). Interestingly, both AS1 and AS2 of the inactive form have similar electrostatic properties: relatively positively charged (II of Fig. 3B) very similar to AS2 of the active form (II of Fig. 2B).

### 2.4. Myricetin destabilizes the active conformation of DengueNS2B-NS3 protease

In the present study, Myricetin has been identified to bind one allosteric site (AS2) in the active form but to two allosteric sites (AS1 and AS2) in the inactive form of Dengue NS2B- NS3 protease. Intriguingly, both AS1 and AS2 have no overlap with the active site. So an immediate question to be answered is how the binding of Myricetin to these sites can lead to the allosteric inhibition of the catalytic activity of Dengue NS2B-NS3 protease. To address this question, we subsequently conducted molecular dynamics (MD) simulations on two Myricetin- protease complexes up to 50 ns. MD simulation represents a powerful method to obtain the dynamic properties of proteins underlying their functions including the enzymatic catalysis (34–38). Particularly, it can pinpoint the dynamically-driven allosteric mechanisms for the enzyme catalysis and inhibition. For example, previously by MD simulations, we have successfully revealed the existence of a global allosteric network in SARS 3C-like protease, the mutation perturbation of which could lead to either inactivation (37) or enhancement of the catalytic activity (38), although all these mutations have no significant effect on the crystal structures. Very recently, with MD simulations we delineated the dynamically-driven mechanism for the allosteric inhibition of Dengue NS2B-NS3 protease by Curcumin (26).

Therefore, here to explore the mechanism by which Myricetin allosterically inhibits the Dengue NS2B-NS3 protease, we first set up three independent MD simulations up to 50 ns on the Myricetin-protease complex in the active form. Fig. 4A presents the structure snapshots in the first set of MD simulations for the active form of Dengue protease in the unbound state (I of Fig. 4A) and in complex with Myricetin at AS2 (II of Fig. 4A). Fig. 4B presents the trajectories of root-mean-square deviations (RMSD) of Cα atoms averaged over three independent simulations of the NS2B cofactor and NS3 protease domain of Dengue protease in the unbound state (blue) we previously performed for understanding the allosteric inhibition of Curcumin (26) as well as in complex with Myricetin we currently conducted (purple). For NS2B, the averaged RMSD values are 5.43 ± 0.52 and 8.09 ± 1.29 Å respectively for the unbound and Myricetin-bound states. For the NS3 protease domain, the averaged RMSD values are 5.58 ± 0.69 and 6.83 ± 0.80 Å respectively for the unbound and Myricetin-bound states. This set of the results clearly indicates that the Myricetin binding led to a large increase of conformational dynamics in NS2B but only a slight change in NS3. This observation is in a general agreement with the NMR results that upon binding to Myricetin, the NS2B factor becomes more dynamic than the NS3 protease domain (Fig. 1C).

**Figure 4.**
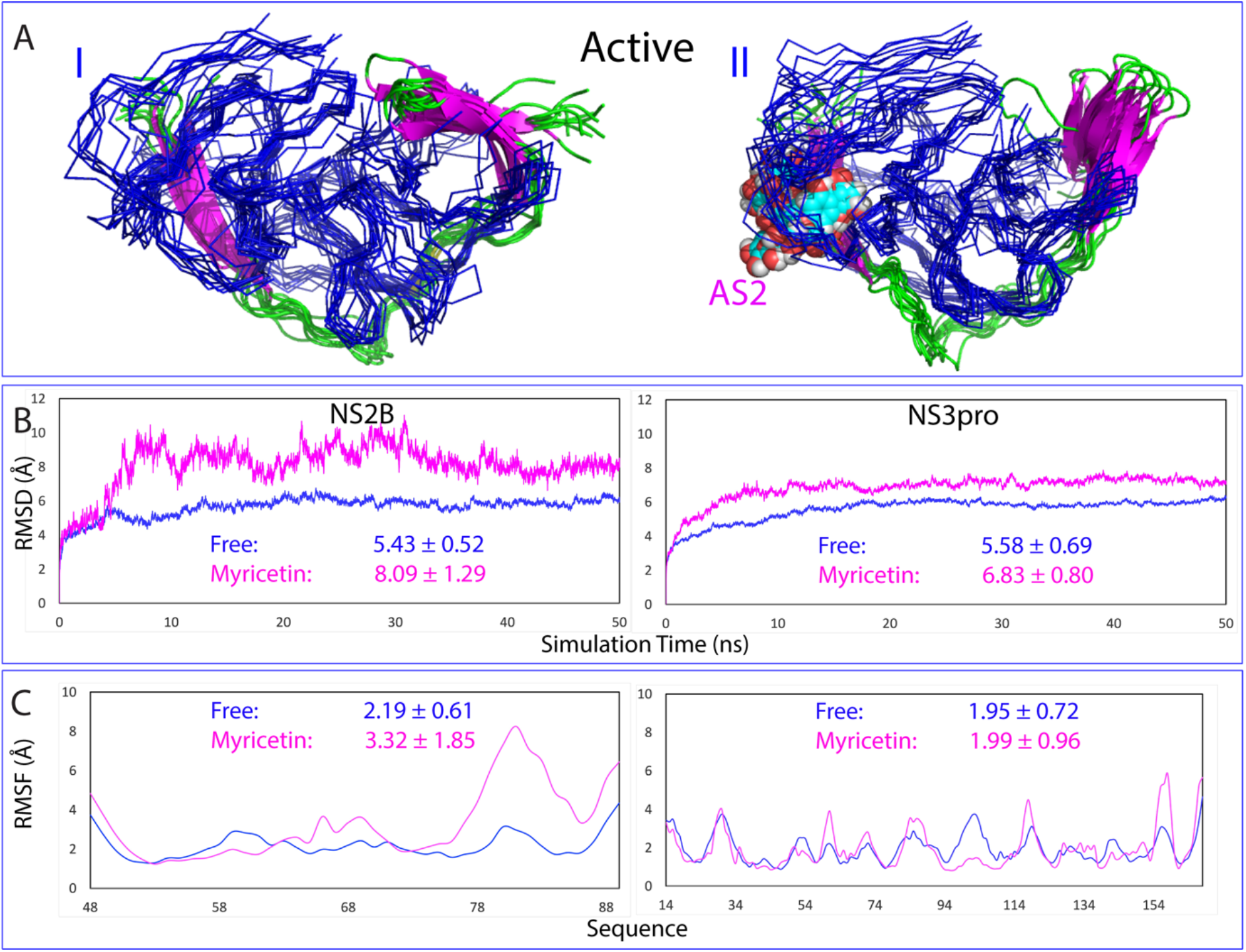
Overall dynamic behaviors of the active form complexed with Myricetin. (A) Superimposition of 11 structures (one structure for 5-ns interval) of the first set of MD simulation for the active Dengue NS2B-NS3 protease in the free state (I), in complex with Myricetin (II). For clarity, NS3 is displayed in the blue ribbon and NS2B in cartoon with the β-strand colored in purple and loop in green; as well as Myricetin in spheres (B) Root-mean- square deviations (RMSD) of the Cα atoms (from their positions in the initial structures used for MD simulations after the energy minimization) averaged over three independent MD simulations of NS2B and NS3 respectively. (C) Root-mean-square fluctuations (RMSF) of the Cα atoms averaged over three independent MD simulations of NS2B and NS3 respectively.

Similar dynamic behaviours are also reflected by the root-mean-square fluctuations (RMSF) of the Cα atoms in the MD simulations. Fig 4C presents the averaged RMSF of three independent simulations, in which for the NS2B residues, the averaged RMSF values are 2.19 ± 0.61 and 3.32 ± 1.85 Å respectively for the unbound and Myricetin-bound states, while for the NS3 residues, the averaged RMSF values are 1.95 ± 0.72 and 1.99 ± 0.96 Å respectively for the unbound and Myricetin-bound states. The RMSF values also indicate that during simulations, no significant difference was observed on the overall conformational fluctuations of the NS3 residues in the unbound and Myricetin-bound states. On the other hand, upon binding to Myricetin, the conformational fluctuations of the NS2B residues becomes increased. Indeed, the C-terminal residues Ile76-Glu89 become highly dynamic, while residues 48-52 and 65-69 also showed high dynamics. On the other hand, it is important to point out that the current 50-ns simulations are insufficient to capture the large conformational transition from the closed (active) to the open (inactive) conformations which is expected to occur over μs-s time scale.

The structural changes in simulations can be better visualized in the structure snapshot at different simulation time points. As well-established, in the active conformation the short antiparallel β-hairpin formed over the NS2B residues Ser75-Ser79 and Gly82-Ile86 wraps the NS3 protease domain to serve as part of the active site pocket and indeed in the previous study its tip was found to have direct contacts with the substrate peptide, which therefore represents a characteristic feature of the active form (Fig. S1A), As we have previously shown (26), although the active form in the unbound state is dynamic, this NS2B β-hairpin retains and only has small fluctuations over its original position in 50-ns simulation. By contrast, for Myricetin- bound state, after 10 ns of the simulation, this short β-hairpin was significantly displaced from its original position to become highly exposed to bulk solvent. Another significant change in the Myricetin-bound state is that the long NS2B β-strand over resides Asp50-Ala57, which have direct contacts with Myricetin, became much shorter after 20 ns, only over Leu51-Glu54 (Fig. 5), while this β-strand showed no significant changes over the 50-ns simulation for the active form in the unbound state (26). Together with our present simulation results on the active form in the unbound Dengue protease (26), the MD simulation results reveal that the binding of Myricetin at AS2 not only disrupt the NS2B β-strand over Asp50-Ala57 which has the direct contacts with Myricetin, but unexpectedly displaced the characteristic β-hairpin from the active site pocket to become highly exposed to bulk solvent. Therefore, the binding of Myricetin to AS2 of the active form is sufficient to disrupt the active conformation, and consequently achieves the allosteric inhibition of Dengue NS2B-NS3 protease.

**Figure 5.**
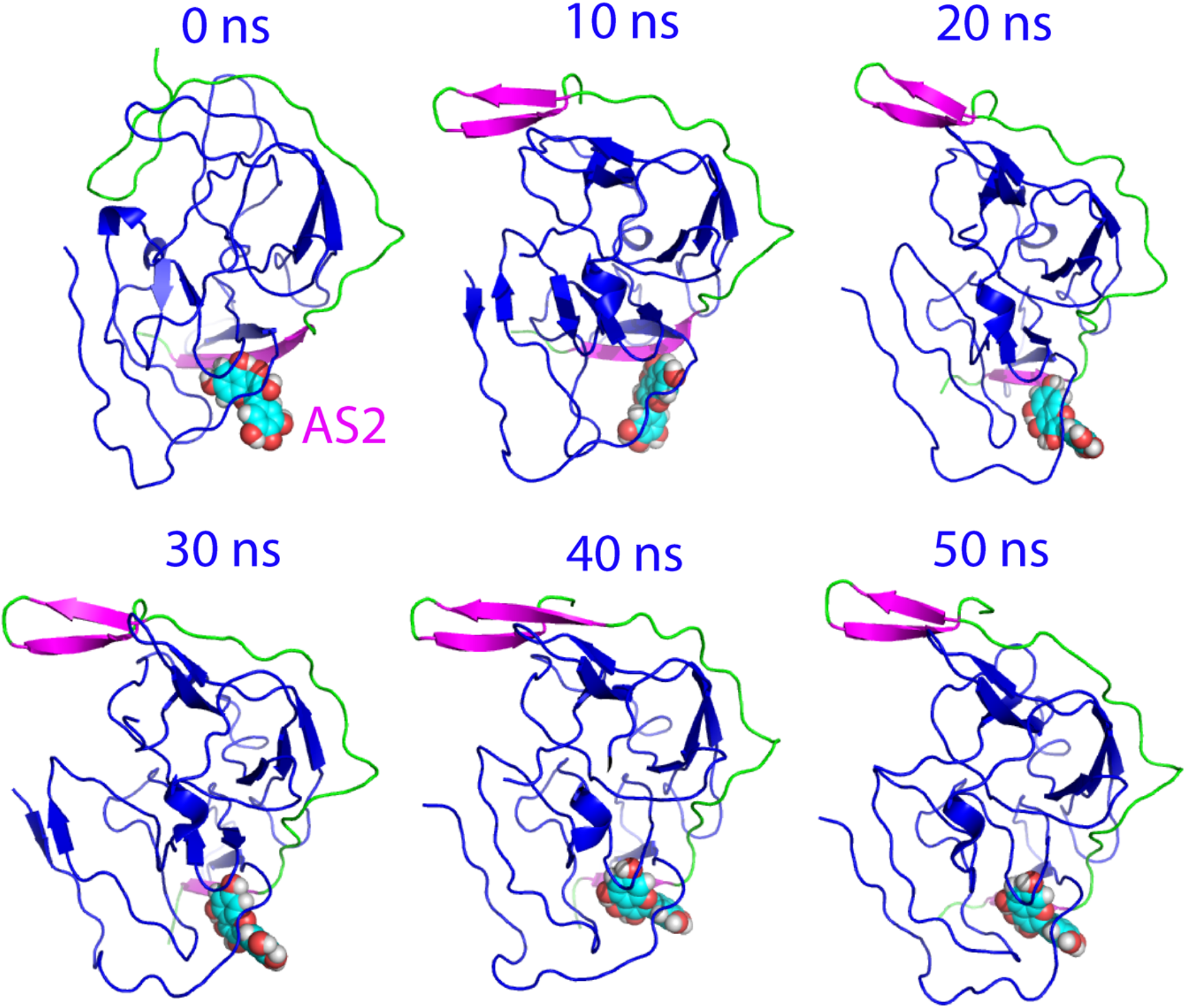
Structure snapshots of MD simulations of the active form. 6 individual structures of the first set of MD simulation for the active form of Dengue NS2B- NS3 protease complexed with Myricetin at different simulation time points. For clarity, the NS3 is displayed in the blue cartoon and NS2B with the β-strand colored in purple and loop in green as well as Myricetin in spheres.

### 2. 5. Myricetin binds and locks the inactive conformation of Dengue NS2B-NS3 protease

We further performed three independent MD simulations up to 50 ns on the inactive form of Dengue NS2B-NS3 protease in the unbound and Myricetin-bound states. Fig. 6A presents the structure snapshots in the first sets of MD simulations in the inactive form in the unbound state (I of Fig. 6A) and in complex with two Myricetin molecules (II of Fig. 6A). Strikingly, overall the inactive form in the unbound state appears to be much less dynamic than that of the active form (Fig. 4A), while the inactive form in the Myricetin-bound state became even less dynamic than the inactive form in the unbound state (Fig. 6A).

**Figure 6.**
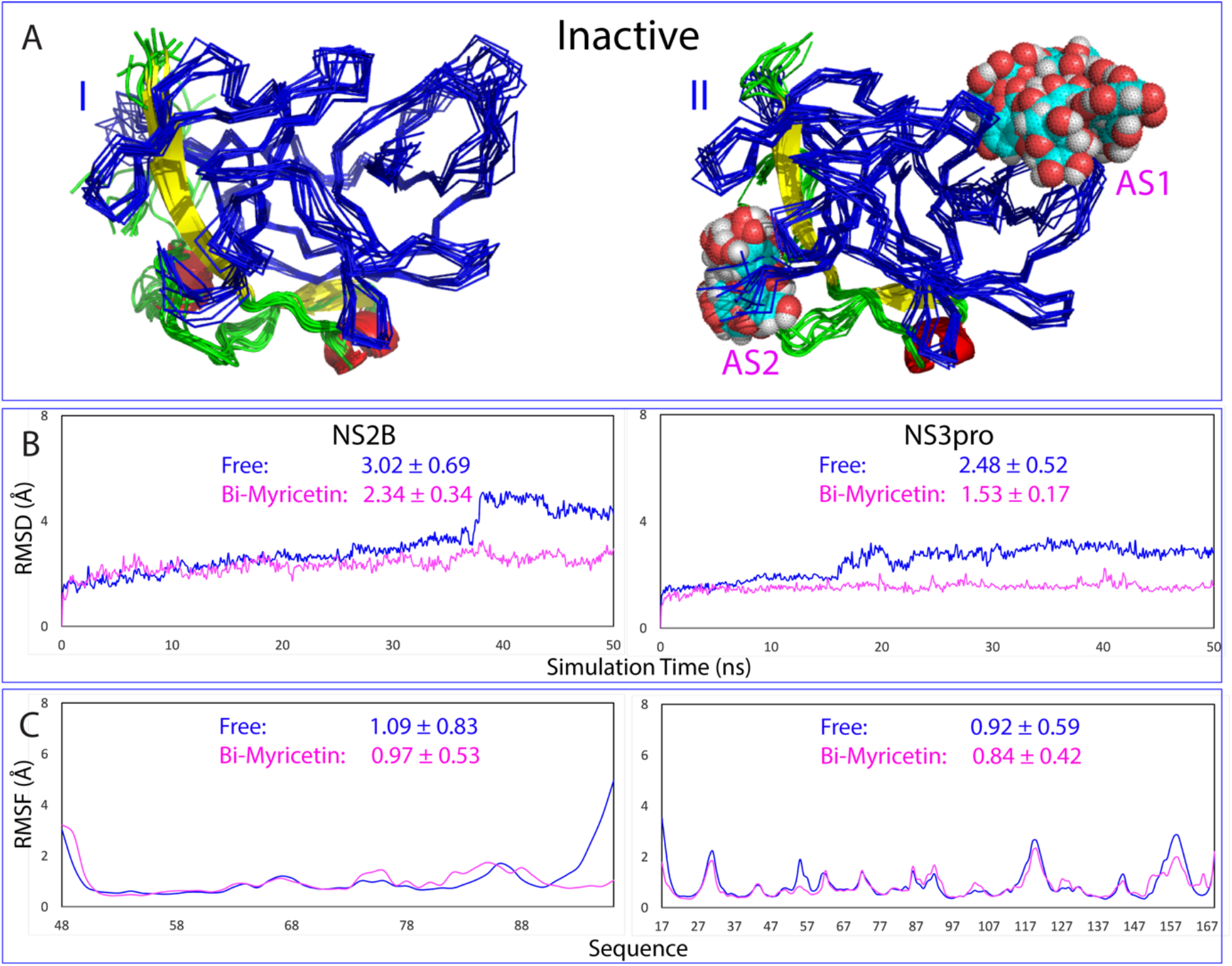
Overall dynamic behaviors of the inactive form complexed with Myricetin. (A) Superimposition of 11 structures (one structure for 5-ns interval) of the first set of MD simulation for the inactive form of Dengue NS2B-NS3 protease in the free state (I), in complex with Myricetin at two allosteric sites (II). For clarity, NS3 is displayed in the blue ribbon and NS2B with the β-strand colored in yellow, helix in red and loop in green; as well as Myricetin in spheres. (B) Root-mean-square deviations (RMSD) of the Cα atoms (from their positions in the initial structures used for MD simulations after the energy minimization) averaged over three independent MD simulations of NS2B and NS3 respectively. (C) Root-mean-square fluctuations (RMSF) of the Cα atoms averaged over three independent MD simulations of NS2B and NS3 respectively.

Fig. 6B presents the trajectories of root-mean-square deviations (RMSD) of Cα atoms averaged over three independent simulations of the NS2B cofactor and NS3 protease domain of the inactive form of Dengue protease in the unbound state (blue) and in complex with two Myricetin molecules (purple). For NS2B, the averaged RMSD values are 3.02 ± 0.69 and 2.34 ± 0.34 Å respectively for the unbound and Myricetin-bound proteases. For the NS3 protease domain, the averaged RMSD values are 2.48 ± 0.52 and 1.53 ± 0.17 Å respectively for the unbound and Myricetin-bound proteases. The results clearly indicate that: 1) even in the unbound state the inactive form is much less dynamic than the active form for both NS2B cofactor and NS3 protease domain. 2) Most strikingly, unlike the active form for which the Myricetin binding provoked the increase in conformational dynamics particularly over the NS2B cofactor, the binding of two Myricetin molecules to AS1 and AS2 of the inactive form by contrast led to the reduction of conformational dynamics for both NS2B and NS3.

Similar dynamic behaviours are also reflected by the root-mean-square fluctuations (RMSF) of the Cα atoms in the MD simulations. Fig 6C presents the averaged RMSF of three independent simulations, in which for NS2B, the averaged RMSF values are 1.09 ± 0.83 and 0.97 ± 0.53 Å respectively for the unbound and Myricetin-bound proteases, while for the NS3 protease domain, the averaged RMSF values are 0.92 ± 0.59 and 0.84 ± 0.42 Å respectively for the unbound and Myricetin-bound proteases. The RMSF values indicate that during simulations, no significant difference was observed on the overall conformational fluctuations of the NS3 residues in the unbound and Myricetin-bound states. On the other hand, upon binding to two Myricetin molecules, the conformational fluctuations of the NS2B residues becomes decreased. Particularly, the RMSF of residues Glu92-Arg100 become significantly reduced (I of Fig. 6C). Moreover, as shown in the structure snapshots (Fig. 7), the inactive forms in both the unbound and bound with two Myricetin molecules are much less dynamic than those of the active forms (Fig. 5).

**Figure 7.**
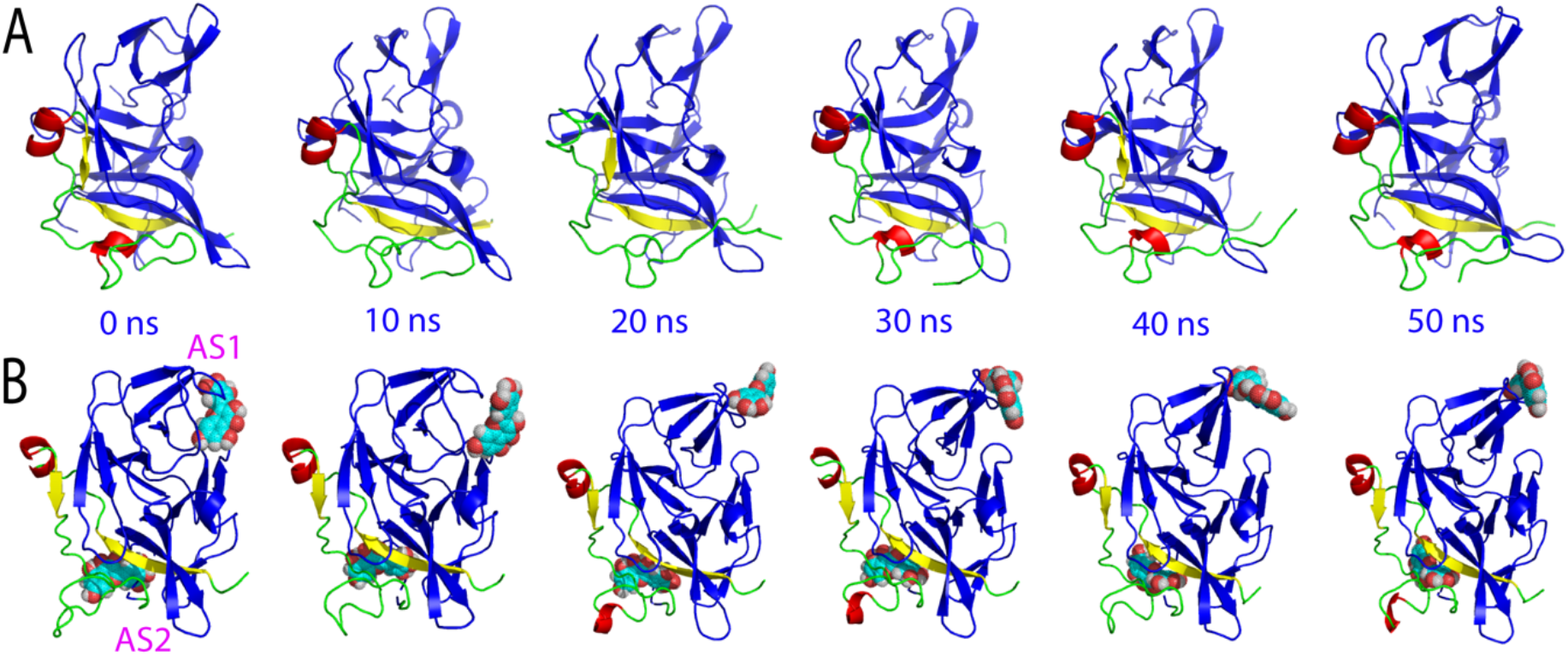
Structure snapshots of MD simulations of the inactive form. (A) 6 individual structures of the first set of MD simulation for the inactive form of Dengue NS2B-NS3 protease in the free state. (B) 6 individual structures of the first set of MD simulation for the inactive form of Dengue NS2B-NS3 protease complexed with Myricetin at two allosteric sites at different simulation time points. For clarity, NS3 is displayed in the blue cartoon and NS2B with the β-strand colored in yellow, helix in red and loop in green as well as Myricetin in spheres.

Therefore, the MD simulation results indicate that the inactive form is less dynamic than the active form and most unexpectedly, the binding of two Myricetin molecules to the inactive form further reduced the conformational dynamics of both the NS2B cofactor and NS3 protease domain. Therefore, Myricetin appears to allosterically inhibit Dengue NS2B-NS3 protease also by binding to both AS1 and AS2 to lock the inactive conformation.

## 3. DISCUSSION AND CONCLUSIONS

The unique NS2B-NS3 proteases are highly conserved in Flaviviruses and have been established as a key target for discovery/design of inhibitors to treat Flavivirus infections (4-24,39-42). Currently, most studies have targeted their active sites for discovery/design of competitive inhibitors (21–23). However, due to the intrinsic features of their active sites, the design of the active-site inhibitors has been demonstrated to be highly challenging. In this context, one solution is to discover/design of their allosteric inhibitors which bind to the cavities other than the active sites. Nevertheless, due to the general challenge in delineating the mechanisms for allosteric processes, only a few attempts were devoted to developing allosteric inhibitors of Flaviviral proteases (17-24,26,39-42). Very recently, by enzymatic assay, NMR characterization and MD simulation, we delineated the mechanism for the allosteric inhibition of Dengue NS2B-NS3 protease by Curcumin, which binds an allosteric site (AS1) close to but having no overlap with the active site pocket (26).

In the present study, we further explored the mechanism for the inhibition of Dengue NS2B-NS3 protease by Myricetin, a flavonoid extensively existing in edible plants with the well-recognised nutraceuticals value (43). Very unexpectedly, although Myricetin and Curcumin both allosterically inhibit Dengue NS2B-NS3 protease, NMR characterization revealed that the pattern of the perturbed NS2B and NS3 residues upon binding to Myricetin is very different from that by binding to Curcumin (26). With NMR-derived constraints, the Myricetin-protease complexes were constructed for both active and inactive forms of Dengue NS2B-NS3 protease. Indeed consistent with NMR results, for the active form Myricetin binds a new allosteric site (AS2) which is far away from both the active site pocket and AS1 for binding Curcumin. By contrast, for the inactive form Myricetin can bind both AS1 and AS2.

So a key question arises as how Myricetin achieves the allosteric inhibition on Dengue NS2B-NS3 proteases if its two binding sites have no overlap with the active site pocket. To answer this question, we conducted MD simulations on both active and inactive forms of Dengue protease in the free state and in complex with Myricetin. Strikingly, for the active form, the binding of Myricetin to AS2 far away from the active site pocket as well as NS2B β-hairpin characteristic of the active form is sufficient to displace this β-hairpin from the original position to become highly exposed to bulk solvent, thus disrupting the active conformation. By contrast, for the inactive form, the binding of Myricetin to AS1 and AS2 reduced the conformational dynamics of both NS2B and NS3, thus acting to lock the Dengue protease in the inactive conformation. In this context, here a speculative mechanism is proposed for Myricetin to allosterically inhibit Dengue NS2B-NS3 protease (Fig. 8). Briefly, as previously established, the active and inactive forms co-exist in solution to reach equilibrium of conformational exchanges on μs-ms time scale, which could be modulated by many factors. For the active form (Fig. 8A), AS2 is accessible and suitable for binding Myricetion, which leads to the significant disruption of the active conformation as reflected by the displacement of the characteristic β- hairpin (Fig. 8B). On the other hand, for the inactive form (Fig. 8C), both AS1 and AS2 become accessible and suitable for binding Myricetin which results in the lock of Dengue NS2B-NS3 protease in the inactive conformation (Fig. 8D). Moreover, it might also be possible that for the active form, upon displacing the β-hairpin by binding of one Myricetin molecule at AS2, AS1 then becomes accessible to binding another Myricetin molecule, which will transform the active form to the inactive form. Unfortunately, as this structural transition is anticipated to occur on μs-ms time scale, it thus remains challenging to be assessed by MD simulations. Nevertheless, the present study reveals that unlike Curcumin which allosterically inhibits Dengue NS2B-NS3 protease only by binding AS1 to destabilize the active conformation, Myricetin allosterically inhibits the enzyme by binding AS2 to destabilize the active conformation as well as by binding both AS1 and AS2 to lock the inactive conformation.

**Figure 8.**
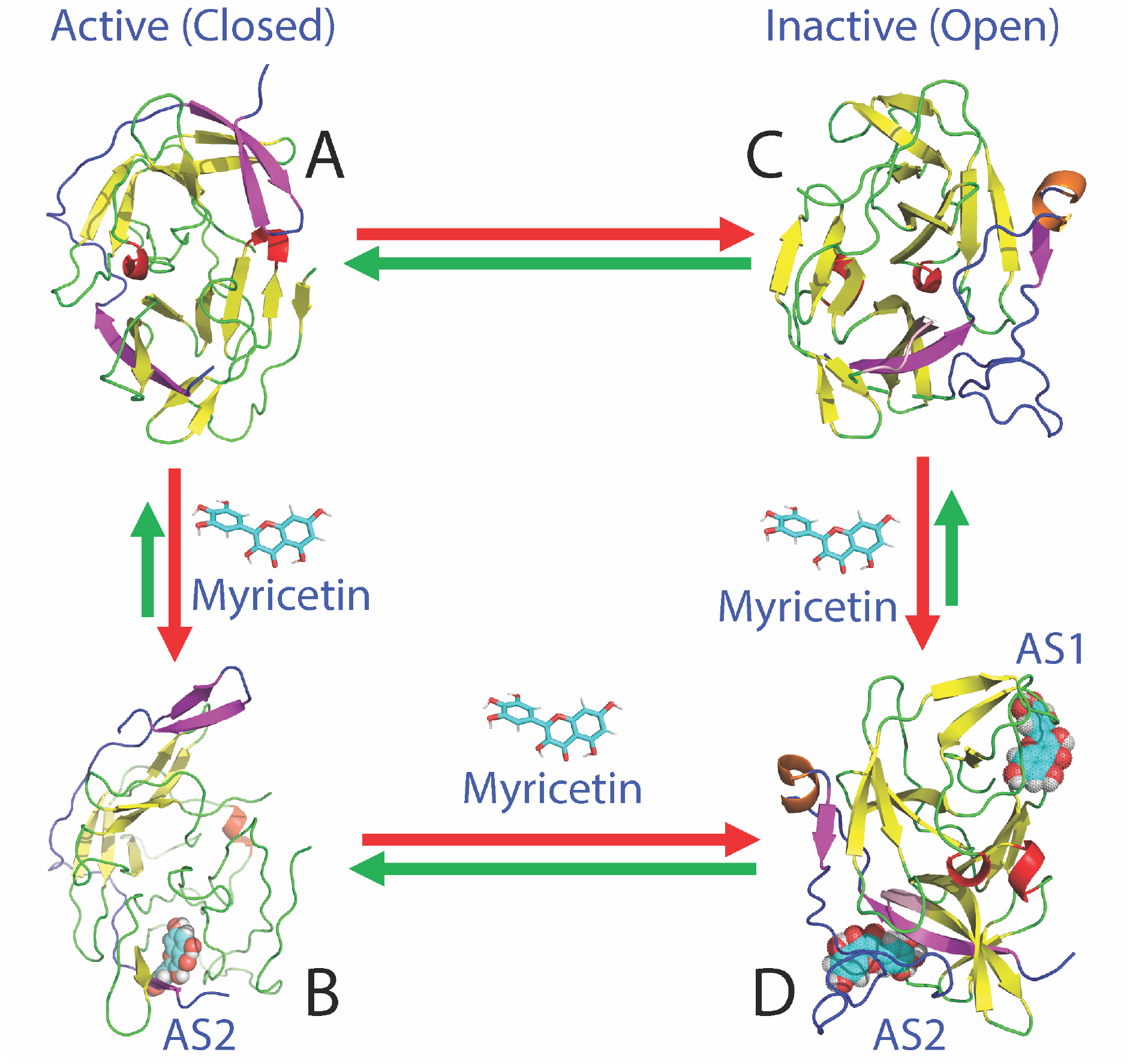
Speculative model for Myricetin to allosterically inhibit Dengue protease. In solution, Dengue NS2B-NS3 protease exists in conformational equilibrium between the active (A) and inactive (C) forms. Myricetin achieves the allosteric inhibition of Dengue NS2B-NS3 protease by binding the active form at AS1 to destabilize the active conformation (B), or/and by binding the inactive form at both AS1 and AS2 to stabilize/lock the inactive conformation (D), both of which lead to the inhibition of the catalytic activity although AS1 and AS2 have no overlap with the active site.

In conclusion, here by NMR and molecular docking, we successfully mapped out the binding modes of Myricetin in both active and inactive forms of Dengue NS2B-NS3 protease. Further MD simulations decode that Myricetin achieves the allosteric inhibition not only by disrupting the active conformation as Curcumin does, but additionally by locking the inactive conformation. With previous results from other and our groups, the current study enforces the notion that Dengue NS2B-NS3 protease has a global allosteric network and perturbation of this network at key sites can achieves allosteric inhibition. The high susceptibility of Dengue NS2B-NS3 protease to the allosteric regulation highlights the promising potential to discover/design its allosteric small molecules of therapeutic application. Finally, given the fact that on the one hand it is extremely daunting to develop a compound into a marketed drug, and on the other hand flavonoids from edible plants including Myricetin have been extensively used for beneficial ingredients of various foods and beverages, Myricetin might be directly utilized to fight the Dengue infections, as well as serves as a promising lead for further design of potent allosteric inhibitors for Dengue NS2B-NS3 protease.

## 4. EXPERIMENTAL SECTION

### 4. 1. Plasmid construction, protein expression and purification

Here we used the expression plasmids we previously constructed (15, 26), which encodes DENV2 NS3 (14–185) and NS2B (48–100) with the same starting and ending residues as the constructs used in the published NMR study (13). The recombinant proteins of NS2B and NS3 were expressed in *Escherichia coli* BL21 (DE3) Star cells following the exactly the same protocol we established before (15, 26). The generation of the isotope-labeled proteins followed the same procedure except for the growth of bacteria in M9 medium with the addition of (^15^NH4)2SO4 for ^15^N labeling and (^15^NH4)2SO4/[^13^C]-glucose for ^15^N-/^13^C-double labelling (15, 26). The purity was checked by SDS-PAGE gels and molecular weights were verified by ESI-MS and Voyager STR time-of-flight-mass spectrometer (Applied Biosystems). The protein concentration was determined by the UV spectroscopic method in the presence of 8 M urea (44).

### 4. 2. Enzymatic kinetic assay

Enzymatic kinetic assay of Dengue NS2B-NS3 protease followed the same protocol we previously described for Dengue and Zika NS2B-NS3 proteases (15,20,26). All enzymatic assays were performed in triplicates. Substrate peptide was Bz-Nle-Lys-Arg-Arg-AMC (Bz-nKRR-AMC), which was purchased from GenScript (Piscataway, NJ), while HPLC-purified Myricetin was from Sigma Aldrich with the purity >95%. Stock solution of Myricetin was dissolved in DMSO. The protease is in 50 mM Tris-HCl (pH 7.5), 0.001% Triton X-100, 0.5 mM EGTA at 37°C. To determine IC50 for Myricetin, 50 nM protease was incubated with various concentrations of Myricetin (in 1μl DMSO) at 37 °C for 30 mins, and Bz-nKRR-AMC addition to 250 µM initiated enzymatic reaction. To determine Ki for Myricetin, the same kinetic assay was performed with different final concentration of Bz-nKRR-AMC and Myricetin. Enzymatic reaction was monitored with fluorescence upon hydrolysis of substrate peptide (Bz-nKRR-AMC) at λex of 380 nm and λem of 450 nm. Fluorescence values (relative fluorescence units/sec) were fitted to the non-competitive inhibition model in GraphPad Prism 7.0; Ki was obtained with fitting to equation: Vmaxinh = Vmax/(1+I/Ki), while I is the concentration of inhibitor (15,20,26). The curves were generated by the program GraphPad Prism 7.0.

### 4. 3. NMR characterization of the binding

2D ^1^H-^15^N HSQC NMR experiments were acquired on an 800 MHz Bruker Avance spectrometer equipped with pulse field gradient units as described previously (15,20,26). NMR samples of 100 μM protease were prepared in 10 mM phosphate buffer, pH 7.5, 5% DMSO and 10% D2O for NMR spin-lock. All NMR experiments were done at 25 °C. Myricetin was also dissolved in the same buffer.

### 4. 4. Molecular docking

In this study, the crystal structures of Dengue NS2B-NS3 protease in the closed form (PDB code: 3U1I) (9) and in the open form (PDB ID of 4M9T) (17) were used for docking. Chemical structure of Myricetin was downloaded from ChemicalBook database (http://www.chemicalbook.com), and their structural geometry were generated and optimized with Avogadro (45). NMR titration derived constraints were used to guide the docking by HADDOCK software (31) and CNS (46). CNS topology and force field parameters of Curcumin is converted from PRODRG server (47). The docking of the Myricetin-protease complexes was performed in three stages: (1) randomization and rigid body docking; (2) semi-flexible simulated annealing; and (3) flexible explicit solvent refinement, as we extensively performed (32, 33). The complex structures with the lowest energy score were selected for the detailed analysis and display by Pymol (PyMOL Molecular Graphics System, Version 0.99rc6 Schrödinger, LLC).

### 4. 5. Molecular dynamics (MD) simulations

The crystal structures of Dengue NS2B-NS3 protease in the active form (PDB code of 3U1I) and in the inactive form (PDB ID of 4M9T) were used as the unbound states, while the docking structures of the Myricetin-protease complexes in the active and inactive forms were used as the bound form for MD simulations with three independent simulations for each of them. Electrostatic potential was first calculated with the 6-31G(d,p) basis set using the GAUSSIAN 16 program, which is converted into partial charge of individual atoms using restrained electrostatic potential (RESP) procedure in Antechamber program (48). The topology parameters of Curcumin were obtained using GAFF (49). All MD simulations reaching 50 ns were conducted using GROMACS (50), and AMBER99SB-IDLN all-atom force field (51) parameters.

The simulation system is a periodic cubic box with a minimum distance of 12 Å between the protein and the box walls to ensure the proteases does not interact with its own periodic images during MD simulation. About 13,000 water molecules (TIP3P model) solvated the cubic box for the all atom MD simulation. 2 Na^+^ ions were randomly placed to neutralize the charge of protein complex. The long-range electrostatic interactions were treated using the fast particle-mesh Ewald summation method (52). The time step was set as 2 fs. All bond lengths including hydrogen atoms were constrained by the LINCS algorithm (53). Prior to MD simulations, the initial structures were relaxed by 500 steps of energy minimization, followed by 100 ps equilibration with a harmonic restraint potential.

## Acknowledgement.

This study is supported by Ministry of Education of Singapore (MOE) Tier 1 Grant R- 154-000-B45-114 and R-154-000-B92-114 to Jianxing Song.

## Supporting Information

The equilibrium of the conformational exchange of Dengue NS2B-NS3 protease in the active and inactive forms, as well as the enzyme of the active form bound with another allosteric inhibitor Curcumin.

## Supplementary Figure

**Figure S1.**
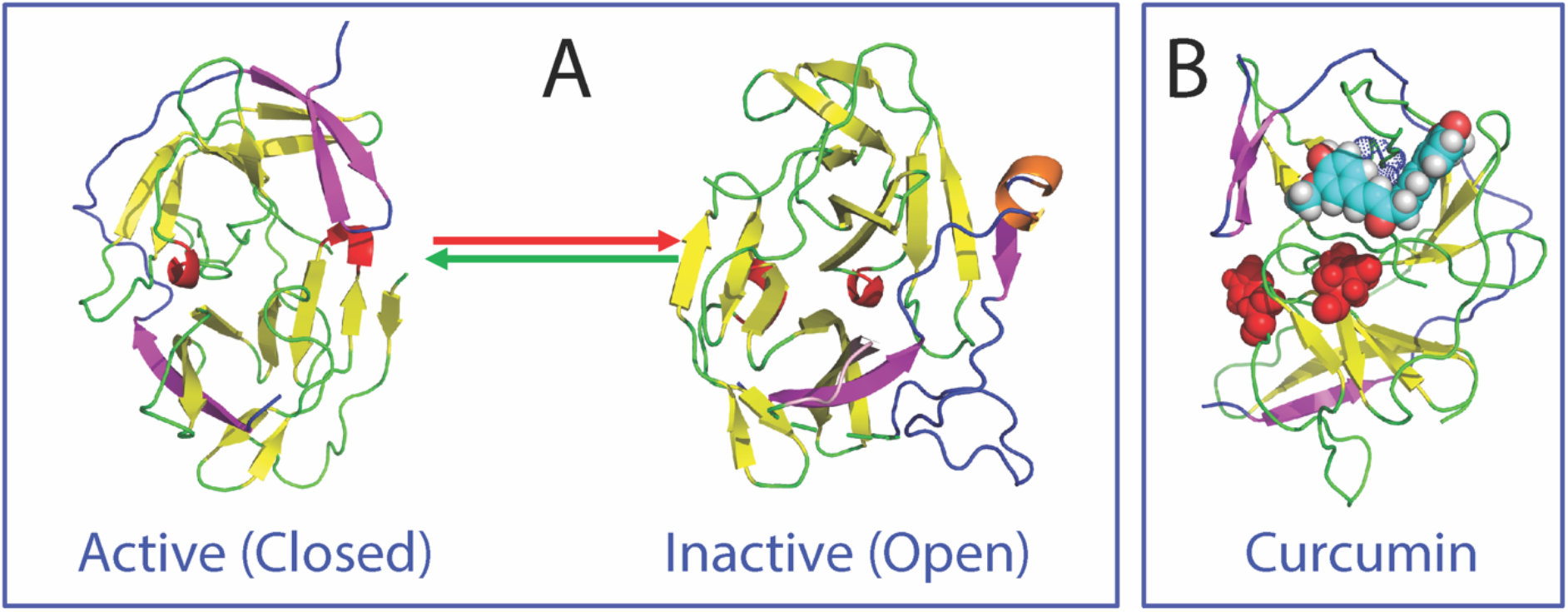
Conformational equilibrium and allosteric inhibition by Curcumin. (A) Conformational equilibrium of the active (closed) and inactive (open) forms of Dengue NS2B-NS3 protease. (B) The active form of Dengue protease in complex with Curcumin (in spheres) at AS1. Red spheres are used to indicate the catalytic triad composed of His51, Asp75 and Ser135.

## Notes

The authors declare no competing financial interest.

### Competing Interest Statement

The authors have declared no competing interest.

